# Spatially Resolved Measurement of Dynamic Glucose Uptake in Live Ex Vivo Tissues

**DOI:** 10.1101/2020.04.17.047068

**Authors:** Austin F. Dunn, Megan A. Catterton, Drake D. Dixon, Rebecca R. Pompano

## Abstract

Highly proliferative cells depend heavily on glycolysis as a source of energy and biological precursor molecules, and glucose uptake is a useful readout of this aspect of metabolic activity. Glucose uptake is commonly quantified by using flow cytometry for cell cultures and positron emission tomography for organs in vivo. However, methods to detect spatiotemporally resolved glucose uptake in intact tissues are far more limited, particularly those that can quantify changes in uptake over time in specific tissue regions and cell types. Using lymph node metabolism as a case study, we developed a novel assay of dynamic and spatially resolved glucose uptake in living tissue by combining ex vivo tissue slice culture with a fluorescent glucose analogue. Live slices of murine lymph node were treated with the glucose analogue 2-[N-(7-nitrobenz-2-oxa-1,3-dia-xol-4-yl)amino]-2-deoxyglucose (2-NBDG). Incubation parameters were optimized to differentiate glucose uptake in activated versus naïve lymphocytes. Regional glucose uptake could be imaged at both the tissue level, by widefield microscopy, and at the cellular level, by confocal microscopy. Furthermore, the assay was readily multiplexed with live immunofluorescence labelling to generate maps of 2-NBDG uptake across tissue regions, revealing highest uptake in T cell-dense regions. The signal was predominantly intracellular and localized to lymphocytes rather than stromal cells. Finally, we demonstrated that the assay was repeatable in the same slices, and imaged the dynamic distribution of glucose uptake in response to ex vivo T cell stimulation for the first time. We anticipate that this assay will serve as a broadly applicable, user-friendly platform to quantify dynamic metabolic activities in complex tissue microenvironments.

## 1. INTRODUCTION

This paper describes a novel, spatially-resolved assay of glycolytic activity in live, intact tissue, and demonstrates its use to quantify dynamic metabolic responses to ex vivo stimulation. Glycolysis is a highly conserved metabolic pathway that converts glucose to lactate and two ATP molecules.^1^ This pathway is strongly upregulated in many types of cells during periods of rapid proliferation, including cancer cells, stem cells, and activated immune cells, even in the presence of oxygen.^2–4^ However, despite the recognition that spatial organization is critical to tissue function,^5–8^ measurement of dynamic glycolytic activity with regional or cellular spatial resolution is still challenging with existing methods.

Uptake of a glucose derivative is a common analytical measure of glycolysis.^9^ In particular, radiolabeled fluorodeoxyglucose (FDG) has a long history for in vivo analysis, and green-fluorescent 2-[N-(7-nitrobenz-2-oxa-1,3-dia-xol-4-yl)amino]-2-deoxyglucose (2-NBDG) has been used more recently for fluorescent detection ex vivo and in vitro.^9,10^ These molecules are only slightly larger than glucose alone and are readily imported through the ubiquitously expressed GLUT receptors for glucose.^11,12^ Studies of 2-NBDG indicate that phosphorylation by hexokinase, the first enzyme in the glycolytic pathway, traps the molecule in the cell (Fig. 1a),^13^ where it remains until it is degraded to a non-fluorescent derivative or dephosphorylated. Thus, the intracellular fluorescent intensity of 2-NBDG provides an estimate of glucose uptake and initial processing that can be detected by flow cytometry or fluorescent microscopy.^9,14,15^

**Figure 1.**
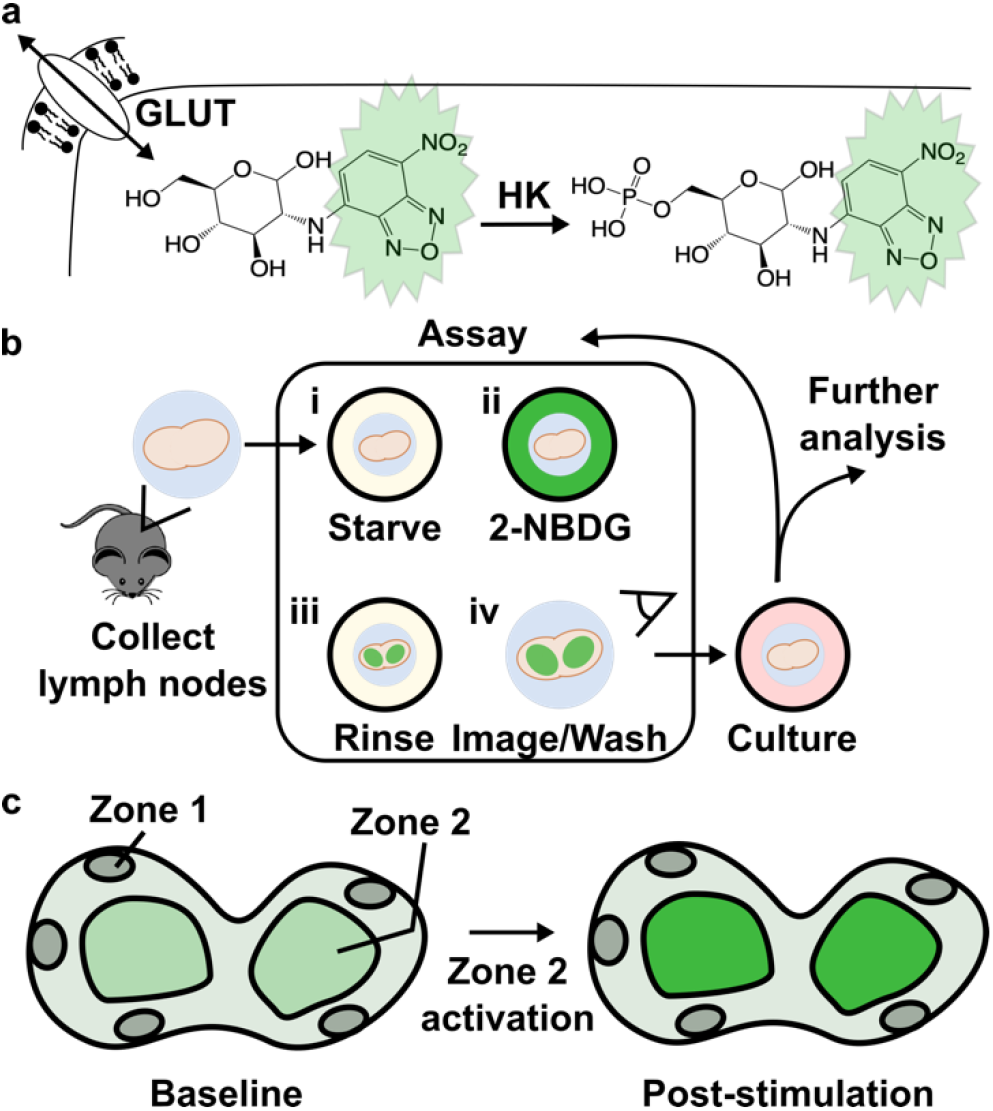
Design of an e*x vivo* metabolic assay to visualize dynamic glucose uptake in live tissue. (a) 2-NBDG is imported through the same GLUT receptors as glucose and temporarily trapped within cells by hexokinase (HK)-mediated phosphorylation. (b) Schematic of assay procedure, which includes a glucose-starvation step, incubation with 2-NBDG and brief rinse, and imaging of the labelled tissue. The assay may be repeated after culture or stimulation, and further analysis such as antibody labeling can be performed after 2-NBDG wash-out. (c) Schematic of a heterogeneous tissue sample, showing hypothetically varied basal 2-NBDG uptake (green shading), and potentially region-specific responses following stimulation.

Even with 2-NBDG and similar reagents, standard assays to measure glucose uptake are limited in their ability to report on spatiotemporal dynamics. Flow cytometric analysis of 2-NBDG uptake in cell suspensions provides high-throughput information at the population level, including after in vitro stimulation, but it provides no insight into the spatial organization of glucose uptake in tissue. In vivo analysis of FDG uptake, on the other hand, detects metabolic activity in situ with spatial resolution on the millimeter scale.^16–19^ This resolution is sufficient for detection of tumors and inflammation, but not to visualize which cells are metabolically active nor how they respond to stimulation. Furthermore, it is difficult to deliver stimuli in vivo at precise concentrations and times, e.g. to observe stimulus-response profiles of glucose uptake.

Ex vivo analysis in live tissue sections offers the potential for both highly resolved imaging and straightforward experimental manipulation. Ex vivo analysis of 2-NBDG uptake has been used primarily for biopsy samples, to identify tumors based on their universally high glucose uptake.^19–22^ These are one-time assays, for which detailed image analysis of uptake at the cellular level is not required and has not been conducted, to our knowledge. Recently, uptake of a Raman-labelled glucose derivative was visualized at the single cell level in mouse brain slices at a single time point.^23^ So far, assays do not exist to measure dynamic glucose uptake over time and with cellular-scale spatial resolution, especially in response to well controlled stimulation.

Interestingly, the nuanced organization of tissue substructures is itself a challenge to testing the impact of stimulation on glucose uptake. In particular, if there is high biological variability in basal glucose uptake between tissue samples, then the large intra-group variability may dwarf small inter-group differences, making it difficult to power experiments adequately.^24^ In this context, performing before-and-after measurements within each sample, e.g., in response to stimulation, sets up each sample as its own control. This approach, called repeated-measures experimental design, greatly improves the statistical power of the experiment and reduces the number of samples required,^25^ although it does require that the assay have minimal effect on tissue health and activity.

Dynamic glucose uptake is particularly interesting in the context of immunology,^26–30^ a system characterized by spatial organization within heterogeneous tissues.^31–34^ The growing field of immunometabolism utilizes measures of metabolic activity to predict immune cell development, differentiation, and activation, and to pinpoint therapeutic targets for immunomodulation.^35–37^ Glucose uptake has been correlated with the activation state of T cells, B cells, and antigen-presenting cells, both in tumor immunology and in the response to vaccination.^38,39^ However, these assays were performed mainly at the cellular level, and it is not yet clear how glycolytic activity is distributed throughout immune tissues such as the lymph node. The lymph node is a highly structured organ, and its sub-structures vary widely in terms of cellular composition and activity.^40,41^ The spatial distribution of glucose uptake in the lymph node is not known in resting nodes nor during an immune response.

Here, we report the first method to quantify dynamic and spatially resolved glucose uptake in live tissue slices, by using 2-NBDG in a repeated measures format (Fig. 1b). The assay was designed to be simple to perform, robust, and compatible with most soft tissues (e.g., brain, lung, tumors). As a proof of principle, we applied 2-NBDG in combination with live immunofluorescent staining to whole, 300 μm-thick slices of live lymph node tissue,^42^ to test both the extent of metabolic heterogeneity and to quantify region-specific responses to ex vivo T cell stimulation (Fig. 1c).

## 2. MATERIALS AND METHODS

### 2.1 Animal care

All animal work was approved by the Institutional Animal Care and Use Committee at the

University of Virginia under protocol #4042, and was conducted in compliance with guidelines from the University of Virginia Animal Care and Use Committee and the Office of Laboratory Animal Welfare at the National Institutes of Health (USA). Male and female C57BL/6 mice were purchased from Jackson Laboratories and used while 6-12 weeks old. Mouse were housed in a vivarium under 12-hour light/dark cycles and given food and water *ad libitum*.

### 2.2 Flow cytometry

Analysis was performed on pooled inguinal, brachial, and axillary lymph nodes from 1 female mouse and 1 male mouse after humane sacrifice via isoflurane overdose and cervical dislocation. Lymph nodes were crushed through a 70-μm filter and centrifuged (Sorvall ST40R Centrifuge, Fisher Scientific) at 400 x g for 5 min. Lymphocytes were cultured at 1 x 10^6^ cells/mL in culture-treated 96-well plates in complete media. Complete media consisted of RPMI 1640 without L-glutamine (Lonza), supplemented with 10% fetal bovine serum (FBS), 1% l-glutamine, 1% Pen-Strep, 50 μM beta-mercaptoethanol, 1 mM pyruvate, 1% non-essential amino acids, and 20 mM HEPES (Fisher Scientific). For stimulation experiments, cells were cultured overnight with or without 4 μg/mL Syrian hamster anti-mouse CD28 (clone 37.51, BioLegend, USA) and plate-bound Armenian hamster anti-mouse CD3ε (clone 145-2C11, BioLegend, USA, coated using 10 μg/mL in PBS overnight at 4°C). Cells were collected, centrifuged, rinsed once in PBS with 2% serum (VWR), and resuspended in a “starve media” (PBS with 10% serum). 2-NBDG (Thermo Fisher) was prepared in 20 mM aliquots in DMSO and stored at −20°C. 2-NBDG was spiked into the starve media at the indicated concentration (25, 100, or 200 μM) and incubated at 37°C and 5% CO_2_ for the indicated time (15, 30, or 45 min). Following 2-NBDG treatment, all cell suspensions were Fc blocked with rat anti-mouse CD16/32 (clone 93, BioLegend, USA) and then stained with 10 μg/mL AlexaFluor647 Armenian hamster anti-mouse CD3ε (clone 145-2C11, BioLegend, USA) followed by 0.5 μg/mL 7-AAD (AAT Bioquest). Single-stain and FMO (fluorescence-minus-one) control conditions were set up as appropriate. For a killed control, 35% EtOH was added to a cell suspension for 20 min at room temperature. Data were collected using a Millipore Sigma Guava easyCyte 6-2L flow cytometer, acquiring data using GuavaSoft 3.3 software.

Compensation and data analysis were performed using FCS Express 6 software. Events were gated on scatter to identify lymphocytes, singlets to identify single cells, and large or CD3+ cells to identify T cells with or without prior activation. Compensation matrices were calculated based on median fluorescent intensity of controls. Median intensity was reported for all samples.

### 2.3 Slice Preparation and Culture

Tissue slicing was performed as previously described.^42,43^ In brief, mice were humanely sacrificed, and inguinal, brachial, and axillary lymph nodes were removed and submerged in ice-cold complete media. Tissues were embedded in 6% low-melting point agarose (Lonza) for two minutes on ice. Agarose blocks were sliced to 300 μm thickness using a vibratome (Leica VT1000S) under sterile conditions. Immediately after slicing, tissue slices were incubated in 1 ml/slice pre-warmed Complete media at 37°C/5% CO_2_ for one hour to recover. Where noted, the media was spiked with phytohaemagglutinin-L (PHA-L) (Millipore Sigma) at a concentration of 25 μg/mL during culture.

### 2.4 Widefield Imaging

For all imaging steps, slices were transferred into a 12-well plate in 0.5 mL PBS for immediate imaging on an upright AxioZoom macroscope using an HXP 200C metal halide lamp, PlanNeoFluor Z 1x/0.25 FWD 56 mm objective, and Axiocam 506 mono camera (Carl Zeiss Microscopy). Filters used were Zeiss Filter Set 38 HE (Ex: 470/40, Em: 525/50), 43 HE (Ex: 550/25, Em: 605/70); 64 HE (Ex: 587/25, Em: 647/70); and 50 HE (Ex: 640/30, Em: 690/50). Brightfield images were also collected for each slice with 10 ms exposure. All fluorescent images of antibody staining were collected at 900 ms. 2-NBDG imaging used 300 ms exposure and filter 38 HE (“FITC”). Zen 2 blue software was used for image collection.

### 2.5 Confocal imaging

After slices had been imaged on widefield microscope, they were immediately returned to ice and transported to a Nikon A1R Confocal microscope. One slice at a time was mounted onto a glass slide and a coverslip was placed on top. A 10x objective was used to locate regions of interest, then slices were imaged with a NIR Apo 40x 0.8W DIC N2 objective. Wavelengths (in nm) of the lasers and filters used were 487 (Emission filter: 525/50), 561 (Emission filter: 595/50), 638 (Emission filter: 700/75). Laser power used for imaging was 0.5, and gain was adjusted for each channel – 95 (FITC), 110 (TRITC), 135 (Cy5). All images were collected between approximately 10-30 μm depth into tissue.

### 2.6 2-NBDG Assay Procedure

A 2-NBDG working solution was prepared fresh for each experiment, consisting of 100 μM 2-NBDG in “starve media” (PBS + 10% FBS). This solution was stored at 4°C when not in use. Staining with 2-NBDG was similar to previously published live lymph node slice staining procedures with other reagents.^44^ First, to record baseline autofluorescence, lymph node slices were rinsed once with 1x PBS and imaged in a tissue culture treated 12-well plate as described above. The PBS was removed and replaced with 900 μL of pre-warmed starve media, and the slices were incubated 30 min (37°C with 5% CO_2_). Next, a 6-well plate was prepared by lining the wells with paraffin film to provide a hydrophobic surface. A 30 μL droplet of the 2-NBDG solution was placed on the parafilm, a tissue slice was carefully transferred on top using a fan brush and weighed down with an A2 stainless steel flat washer (10 mm outer diameter, 5.3 mm inner; Grainger, USA), and another 30 μL droplet of 2-NBDG was added on top to ensure staining on both the sides of the tissue. The 6-well plate was covered and incubated for 30 min (37 °C, 5 % CO_2_). Next, slices were rinsed by soaking in 1.5 mL 1x PBS per slice for 15 min (4 °C), shaking gently halfway through the rinsing period. Slices were transferred to 0.5 mL chilled 1x PBS and immediately imaged. Slices were returned to complete media and incubated (37 °C, 5 % CO_2_) pending further use. Care was taken not to flip the slices over at any point during the assay procedure.

### 2.7 Immunostaining of live lymph node slices

Live fluorescence immunostaining was performed as previously reported.^44^ Briefly, live slices were transferred placed on parafilm, Fc-blocked for 30 min, and labelled with an antibody cocktail for one hour. Antibodies used were FITC-rat anti-mouse/human CD45R/B220 (clone RA3-6B2, BioLegend), and rat anti-mouse CD4 F(ab’)_2_ fragment (clone GK1.5, BioLegend, fragmented according to published procedures^45^) that was conjugated using AlexaFluor 594-succinimidyl ester according to manufacturer instructions (Thermo Fisher). All antibodies were used at a concentration of 20 μg/mL (0.2 μg per tissue slice). Slices were rinsed for 1 hr in PBS to remove unbound antibodies.

### 2.8 Image Analysis

Images of slices were analyzed using Fiji v1.48.^46^ T cell rich regions-of-interest (T cell zone ROI; CD4^+^B220^-^) were identified manually from immunofluorescence images, by a researcher who was blinded to the images with 2-NBDG. Images of 2-NBDG were then rotated and translated to match the orientation of the tissue in the antibody stained images, and the traced regions of interest were copied onto them. The mean fluorescent intensity (MFI) of 2-NBDG signal was quantified in the T cell zone ROI and, separately, in the areas of tissue excluded from that ROI (“non-T cell zone”). Where noted, the entire slice was traced instead, by identifying the border based on brightfield images. 2-NBDG intensity (2-NBDG Int.) values are reported with autofluorescence subtracted out, as follows,

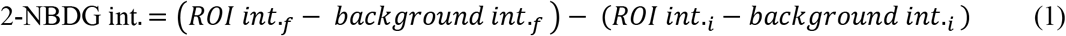

where “*i*” (initial) represents the intensity prior to 2-NBDG treatment (i.e., autofluorescence) and “*f*” (final) represents the intensity measured after the assay with 2-NBDG. “Background int.” for each image was defined as the average of three regions of the image not containing tissue. Slices that had little T cell zone, defined as fewer than 1.5×10^5^ μm^2^ contiguous area, were excluded from analysis. For 2-NBDG image display, brightness and contrast were adjusted uniformly across all slices that are displayed in the same figure.

### 2.9 Statistics

Statistical tests and curve fits were performed in GraphPad Prism 8.3.0.

## 3. RESULTS AND DISCUSSION

### 3.1 Optimization of 2-NBDG treatment to detect T cell activation in mixed cultures

Our goal was to develop an assay that could detect the increased glucose uptake in a lymph node after immune activation.^26^ In preliminary experiments, application of 2-NBDG to murine lymph node slices under unoptimized conditions (2-NBDG concentration and time; rinse time) was unsuccessful. There was little difference in 2-NBDG intensity or distribution between live and ethanol-killed samples (data not shown), indicating most of the signal did not arise from metabolic processes.^14^ Therefore, we sought to identify assay conditions that maximized the sensitivity to 2-NBDG uptake due to active metabolism, not just passive diffusion or non-specific binding.^47^

To distinguish activated from resting lymphocytes, we optimized both the concentration and time of incubation with 2-NBDG by flow cytometry, as has been done previously for other cell types.^15,48–50^ Mixed murine lymphocytes were cultured overnight with or without T cell stimulation (anti-CD3 and anti-CD28). Subsequent 2-NBDG treatment was conducted in a “starve media” consisting of PBS containing 10% FBS.^51^ This media provided some minimal nutrients and proteins, while minimizing the amount of D-glucose, which competes with 2-NBDG for receptor binding.^13^ To identify resting and activated T cells by flow cytometry, the lymphocyte population was gated on live cells that were CD3-positive and/or expanded in size (high forward scatter) (Fig. 2a-c). We note that this gating strategy, necessitated by the internalization of CD3 after anti-CD3 stimulation,^52^ may capture any large CD3-negative cells as well.

**Figure 2.**
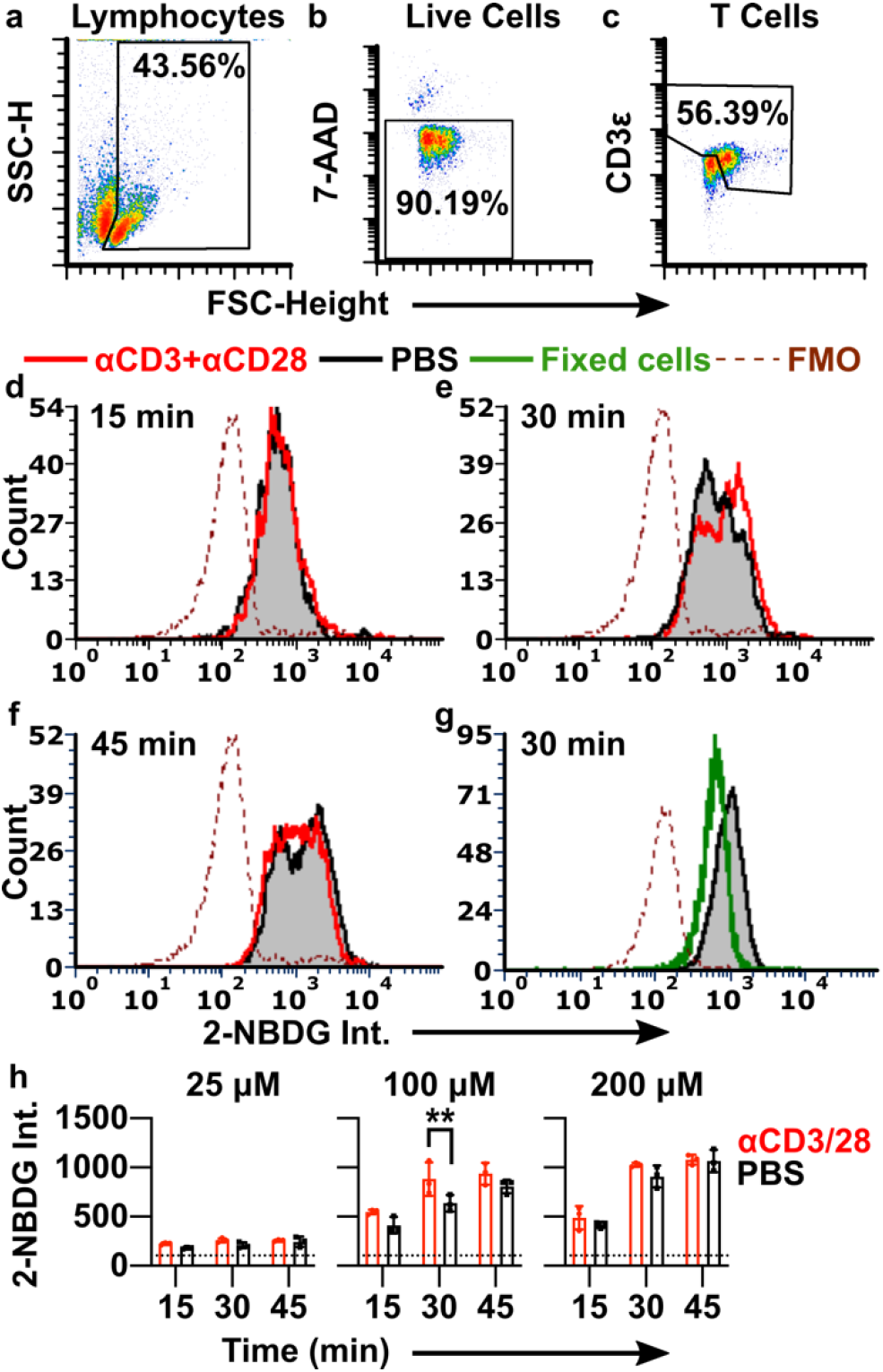
Optimization of 2-NBDG labelling conditions to detect T cell activation. (a-c) Events were gated on forward and side scatter to identify lymphocytes (a), singlets to identify single cells (not shown), 7-AAD exclusion to identify live cells (b), and a combination of high expanded forward scatter and CD3 expression to identify resting and activated T cells (c). SSC-H shown on linear scale; 7-AAD and CD3 intensities shown on log scale. (d-g) Histograms of 2-NBDG intensity after labelling with 100 μM 2-NBDG for 15 min, 30 min, or 45 min. Lymphocytes were stimulated with anti-CD3 and anti-CD28 (red) or left unstimulated (black/grey shaded). The dotted line is the fluorescence-minus-one (FMO) control for 2-NBDG. Histograms were normalized to 5000 events. (g) 2-NBDG labeling of live and formalin-fixed cells after 30 min, 100 μM 2-NBDG. Fixed samples were gated on singlet events. (h) Median 2-NBDG intensity depended on incubation time and concentration of 2-NBDG. Bars show mean ± std dev of N=3 replicates. Dotted lines show mean of FMO (no 2-NBDG) control. Two-way ANOVA with Sidak’s multiple comparisons, ** represents p<0.01.

The ability to distinguish activated from naïve T cells depended strongly on the incubation conditions (Fig. 2e-h). Even formalin-fixed lymphocytes were labelled to some extent by 2-NBDG, suggesting that small amounts of 2-NBDG diffused passively into or adhered to the surface of non-viable cells (Fig. 2g). After a short, 15-min 2-NBDG treatment (100 μM), live cells showed a single peak that was indistinguishable between naïve and stimulated conditions (Fig. 2d), and the signal often overlapped with that of fixed controls (not shown). These data suggest that the signal after a 15 min incubation was due to passive labelling rather than quantifiable metabolic activity. In contrast, after a 30-min 2-NBDG treatment, a new peak appeared in the stimulated condition, creating a characteristic “doublet” pattern with two closely overlapping peaks (Fig. 2e). The 1.3-fold difference in intensity between the two peaks, though small, was consistent with prior reports showing a 1.3 – 2.0-fold increase in 2-NBDG uptake by T cells after metabolic stimulation (Fig. 2h).^27,49^ We concluded that the brighter peak represented increased retention of 2-NBDG driven by stimulation-induced metabolism. In a separate experiment, inclusion of high D-glucose concentrations (11 mM in RPMI versus ~700 μM in the starve media) during 2-NBDG treatment prevented the appearance of the brighter peak, consistent with the requirement for uptake via glucose receptors expected in mammalian cells (Fig. S1).^13,14^ For reference, physiological blood glucose levels average 9.1 mM in C57Bl/6 mice.^53^ Finally, in agreement with reports in kidney and heart cells,^54,55^ we observed evidence of saturation of 2-NBDG uptake at longer times (45 min) (Fig. 2f) and higher concentrations (200 μM) (Fig. 2h). Under each of these conditions, even naïve cells featured the doublet pattern and the median fluorescent intensity levelled off. Lower concentrations of 2-NBDG (25 μM) produced only low-intensity singlet peaks, with no detectable effect of stimulation (Fig. 2h). As the response to stimulation was best detectable after 30 min staining with 100 μM 2-NBDG, this condition was selected as the appropriate condition to test in lymphoid tissue.

### 3.2 Establishing conditions for 2-NBDG labeling and repeated analysis in live tissue

The overall assay procedure for tissue slices consisted of pre-imaging the tissue to record autofluorescence, followed by a short incubation in starve media, immersion in 2-NBDG solution, a rinse in PBS to remove unbound reagent, and imaging once more. Unlike analysis in cell culture, analysis of glucose uptake in thick slices of tissue required consideration of tissue-specific properties: restricted mass transport (slow diffusion) that may impede rinsing of excess 2-NBDG, potential for non-specific binding to the matrix, and tissue heterogeneity and autofluorescence. We addressed each of these issues to validate the robustness of the ex vivo assay.

To confirm that the majority of signal in live tissues was not due to residual extracellular signal or non-specific matrix binding, labelled slices (100 μM 2-NBDG, 30 min incubation, 15 min rinse) were imaged by confocal microscopy. The 2-NBDG signal appeared primarily intracellular, with well-defined bright circular regions consistent with the dimensions of lymphocytes (Fig. 3a,b). The darkness of the background compared to unstained slices suggested that extracellular 2-NBDG had largely diffused out during the 15-min rinse. These data agreed with theoretical estimates for the diffusion time, *t* [s], of 2-NBDG (*t* = *x*^2^ / 2*D*, where *x* is distance [cm] and *D* is the diffusion coefficient [cm^2^/s]). Using the diffusion coefficients of glucose in brain and eye as a rough estimate (*D* = ~0.91 x 10^-6^ - 2.6 x 10^-6^ cm^2^/s),^56^ extracellular 2-NBDG should diffuse through a 300 μm thick tissue slice in ~3-8 min, significantly less than the 15-min rinse time.

**Figure 3.**
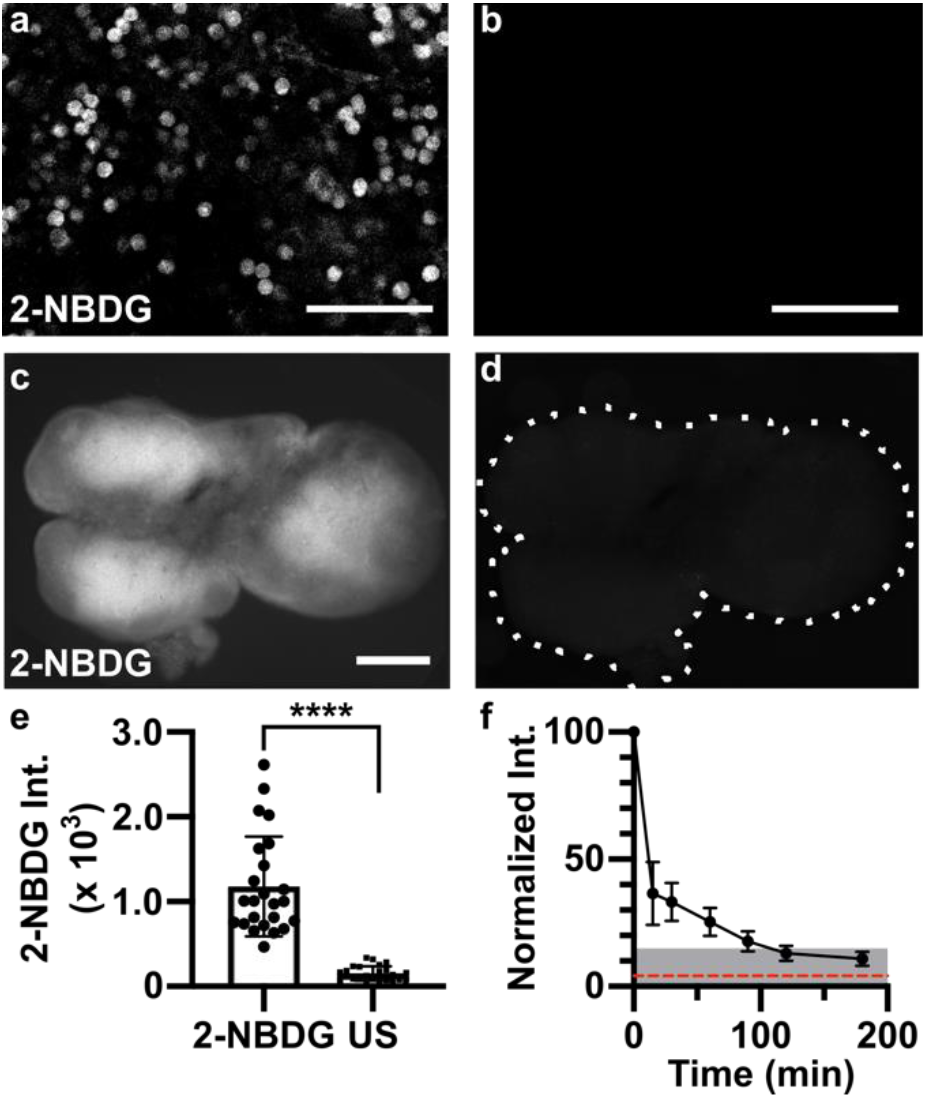
Applying 2-NBDG for quantitative analysis in live tissue. (a,b) Representative confocal images in live slices of naïve lymph node, for (a) a slice incubated with 2-NBDG (30 min, 100 μM) and rinsed for 15 min, and (b) a separate unstained slice, showing minimal autofluorescence (same brightness and contrast). Tissues imaged at a depth of 20-30 μm. Scale bars 50 μm. (c,d) Representative widefield images in a live naïve lymph node slice, (c) after and (d) prior to 2-NBDG labelling (same brightness and contrast). Scale bar 500 μm. Dotted line traces slice border. (e) Mean 2-NBDG intensity in naïve slices with or without 2-NBDG staining (US, unstained), from widefield imaging. Bars show mean ± standard deviation, N=24. **** represents p<0.0001 by paired t-test. (f) Normalized 2-NBDG intensity, measured over time as slices were incubated in media after a single staining step (N=7 slices). Mean intensity data from each slice were normalized to the intensity of that slice at t=0 min. Dotted red line shows mean intensity of unstained slices. Shaded area represents less than 15% of original signal remaining.

Next, we considered tissue heterogeneity and autofluorescence. Unlike well mixed cell suspensions, tissue slices are likely to contain significant heterogeneity in terms of cellular population, location, metabolic state, and autofluorescence, both within and between samples. Indeed, widefield fluorescence imaging revealed clearly defined regions of high and low 2-NBDG uptake across naïve lymph node slices (Fig. 3c), which were clearly distinguishable from tissue autofluorescence (Fig. 3d). In addition, the mean 2-NBDG intensity varied by up to 5-fold between lymph node slices (Fig. 3e), consistent with the fact that each tissue slice contains a unique combination and number of lymphocytes and other cell types, likely at varied density and with varied permeability of the extracellular matrix.^42^ This variability, coupled with the expected small change in 2-NBDG uptake upon stimulation (< 1.5-fold, Figure 2), led us to the conclusion that a repeated-measures experimental design, in which the same tissue would be measured before and after stimulation, would be far more powerful than comparing separate groups of stimulated versus unstimulated slices.

To enable repeated analysis of the same tissue slice, we characterized the minimum recovery time needed for the 2-NBDG signal to decay (Fig. 3f). Based on prior reports, decay of cellular 2-NBDG signal over time is due to a combination of dephosphorylation and export of 2-NBDG and intracellular decomposition into a non-fluorescent form.^11^ After 30-min incubation with 2-NBDG and a 15-min rinse, tissues were cultured in complete media at 37 °C, 5 % CO_2_ and imaged periodically. The 2-NBDG signal diminished by >85% after 2 hr. Based on these results, in future experiments we waited at least 2 hr before repeating the assay, and collected a new “initial” image immediately prior to each 2-NBDG treatment to ensure that any residual 2-NBDG was properly subtracted (Eq. 1).

### 3.3 Mapping spatially resolved 2-NBDG uptake in tissue

To interpret the spatial distribution of 2-NBDG uptake in lymph node slices, we followed the 2-NBDG assay with live immunofluorescence labelling to identify two regions of interest – the deep paracortex, also referred to as the T cell zone (CD4^high^ B220^low^), and the remainder, which includes B cell follicles (B220^high^ circular zones) and sinuses (B220^med^) (Fig. 4a).^44^ We observed that in naïve tissues, T cell zones appeared significantly brighter with 2-NBDG than the small B cell follicles or sinus regions (Fig. 4b-d, Fig. S2). In fact, B cell follicles were often clearly apparent as darkened patches on the slice periphery in the 2-NBDG image. B cell follicles, like the rest of the lymph node, are densely packed with cells, suggesting that cell density was not the driving force behind the difference in glucose uptake.^57^

**Figure 4.**
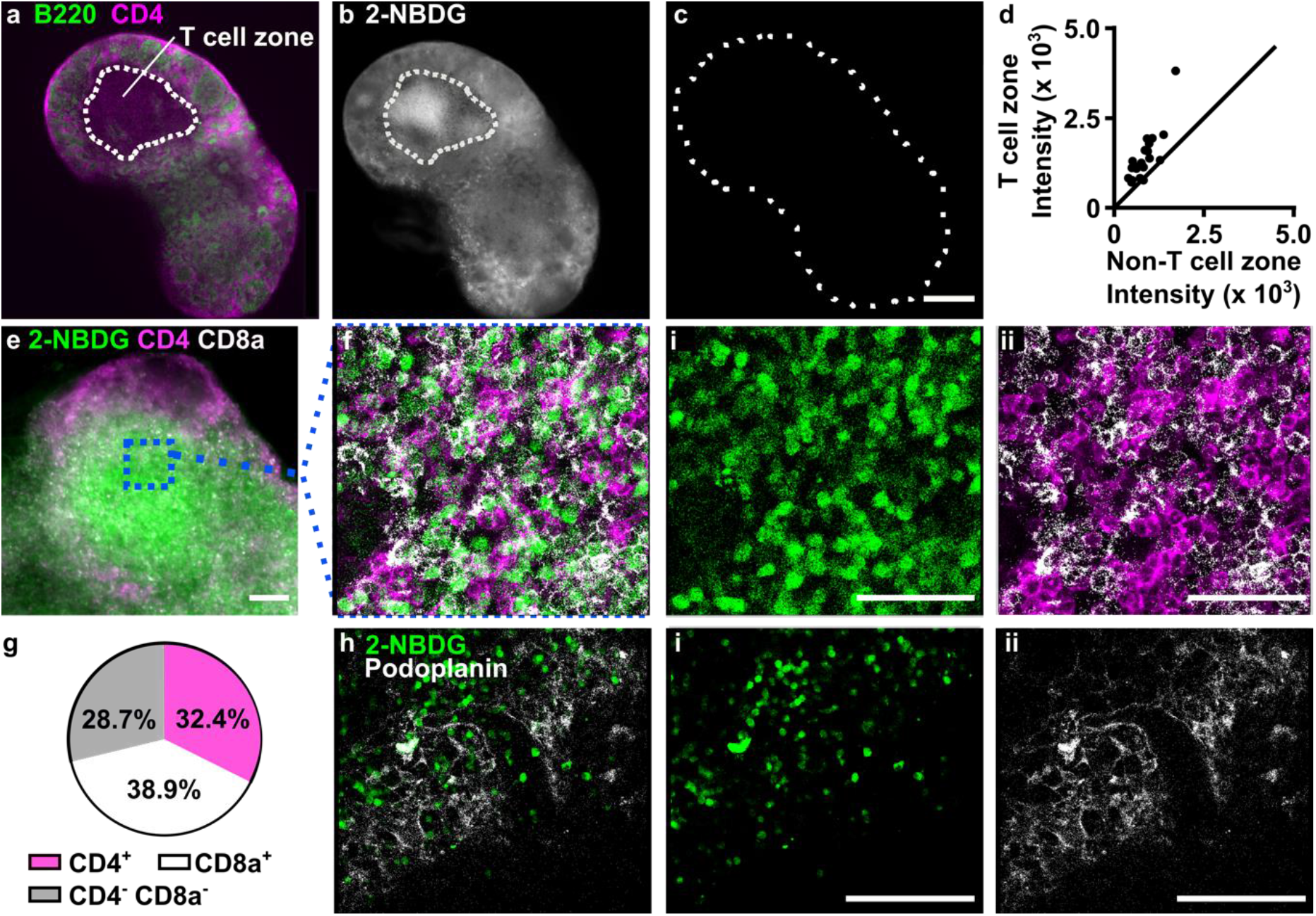
Spatial distribution of 2-NBDG uptake coupled with immunofluorescence. (a) Representative widefield immunofluorescence image of a naïve lymph node slice. Outline represents blinded tracing of T cell zone (CD4^+^B220^-^ area). (b-c) Images of 2-NBDG, collected (b) after 2-NBDG staining and (c) prior to staining in the same slice. Dotted line shows slice border. 2-NBDG images scaled to same brightness and contrast. Scale bar 500 μm. (d) Region-specific mean 2-NBDG intensity within naïve lymph node slices. Intensity was measured inside and outside of T cell zones (CD4^+^B220^-^). The line y = x was plotted for reference. Each dot represents one slice. (e) Representative widefield image of a lymph node slice after live immunostaining and 2-NBDG assay. Scale bar 100 μm. (f) Confocal imaging of the boxed region in panel e, shown as three-color overlay and with separate channels for (i) 2-NBDG uptake and (ii) CD4 (pink) and CD8a (white). Scale bars 50 μm. (g) Manual cell counts determined the percentage of 2-NBDG-positive cells in the T cell zones that were co-stained with CD4, CD8a, or neither. Data compiled from N=11 tissue slices from three mice. (h) Image of 2-NBDG uptake overlaid with immunofluorescence for podoplanin, a surface marker for lymphoid stroma, and separate channels for (i) 2-NBDG uptake and (ii) podoplanin. Representative of N=7 slices from two mice. Scale bars 100 μm.

The bright signal observed in the T cell zone could be due to T cells (CD4^+^ or CD8^+^), the supporting stromal fibroblasts (podoplanin-expressing fibroblastic reticular cells, FRCs), or the less abundant antigen presenting cells. To determine which cell types contributed the most to the 2-NBDG signal in the T cell zone, the lymph node slice was pre-labelled with fluorescent antibodies for cell surface markers, and treated with 2-NBDG immediately afterwards. This imaging strategy allowed for immunofluorescence and glucose uptake to be analyzed simultaneously, avoiding the need for repeated imaging of the tissue. Central regions high in 2-NBDG uptake were identified by widefield imaging and then analyzed at the cellular level by confocal microscopy (Fig. 4e,f). Cell surface staining of CD4 and CD8 formed roughly circular outlines, some of which bounded a circular 2-NBDG bright region, and some of which were dark inside (no detectable 2-NBDG uptake). Of the 2-NBDG-bright regions, most (71%) were surrounded by staining for either CD4 or CD8 (Fig. 4g), suggesting that they were T cells. We considered the possibility that the remaining 2-NBDG-high cells (CD4^-^ CD8^-^) could be a stromal population. However, the FRC marker podoplanin showed minimal overlap with 2-NBDG (Fig. 4h), indicating that these cells did not contribute to the 2-NBDG positive population in the slices. The identity of the 2-NBDG^+^ CD4^-^ CD8^-^ cells was not further investigated here. In summary, these experiments showed that simple incubation with 2-NBDG could reveal spatially heterogeneous glucose uptake in live tissues, and that coupling with live immunofluorescence could provide phenotyping of glucose-rich regions at both the whole-tissue and cellular length scales.

### 3.4 Quantifying local changes in 2-NBDG uptake in response to ex vivo stimulation

Having shown that a single round of the 2-NBDG assay could map out the distribution of glucose uptake, we next addressed the challenge of quantifying responses to ex vivo stimulation over time. As discussed above, both cell compositions and basal metabolic rates vary widely between tissue slices (Fig. 3e), and this variance could mask small changes in glucose uptake between stimulated and unstimulated tissues. We reasoned that the effects of phenotypic variability could be minimized by using a repeated measures experimental design.

To test this hypothesis, we performed the 2-NBDG assay repeatedly in single tissue slices, before and after a metabolic activator was applied. In this manner, each slice served as its own control, thus accounting for the variation in basal metabolic activity within and between slices. Phytohemagglutinin (PHA) was used as a model immunostimulatory treatment. PHA crosslinks glycosylated surface receptors such as the T cell receptor (TCR), leading to upregulation of glycolysis within 3 hours, particularly in the T cell population.^58–61^ Murine T cells stimulated via the TCR upregulate surface GLUT1 receptors within 2-4 hr, an early hallmark of T cell activation.^62^ Additionally, increased hexokinase activity has been observed after 2 hr PHA stimulation of mixed lymphocytes.^63^ We therefore hypothesized that a 3-hour culture with PHA would increase 2-NBDG uptake primarily in the T cell zone. Autofluorescence and basal 2-NBDG uptake were assayed as above, and then the lymph node slices were cultured with or without PHA for 3 hours. In addition to stimulation, this period also provided sufficient time clear the majority of the 2-NBDG signal from the first round of the assay (Fig. 3f). Afterwards, the 2-NBDG assay was repeated, and finally the slices were immunostained for B220 and CD4 to define the tissue architecture. The immunostaining step was performed last to avoid antibody-mediated T cell activation during the multi-hour culture period.^44^ As before, basal 2-NBDG uptake was greatest in the T cell zone (CD4^high^ B220^low^ regions), with the remaining regions – a composite of B cell follicles, sinuses, and macrophage-rich medulla – remaining dark (Fig. 5, Fig. S2).

**Figure 5.**
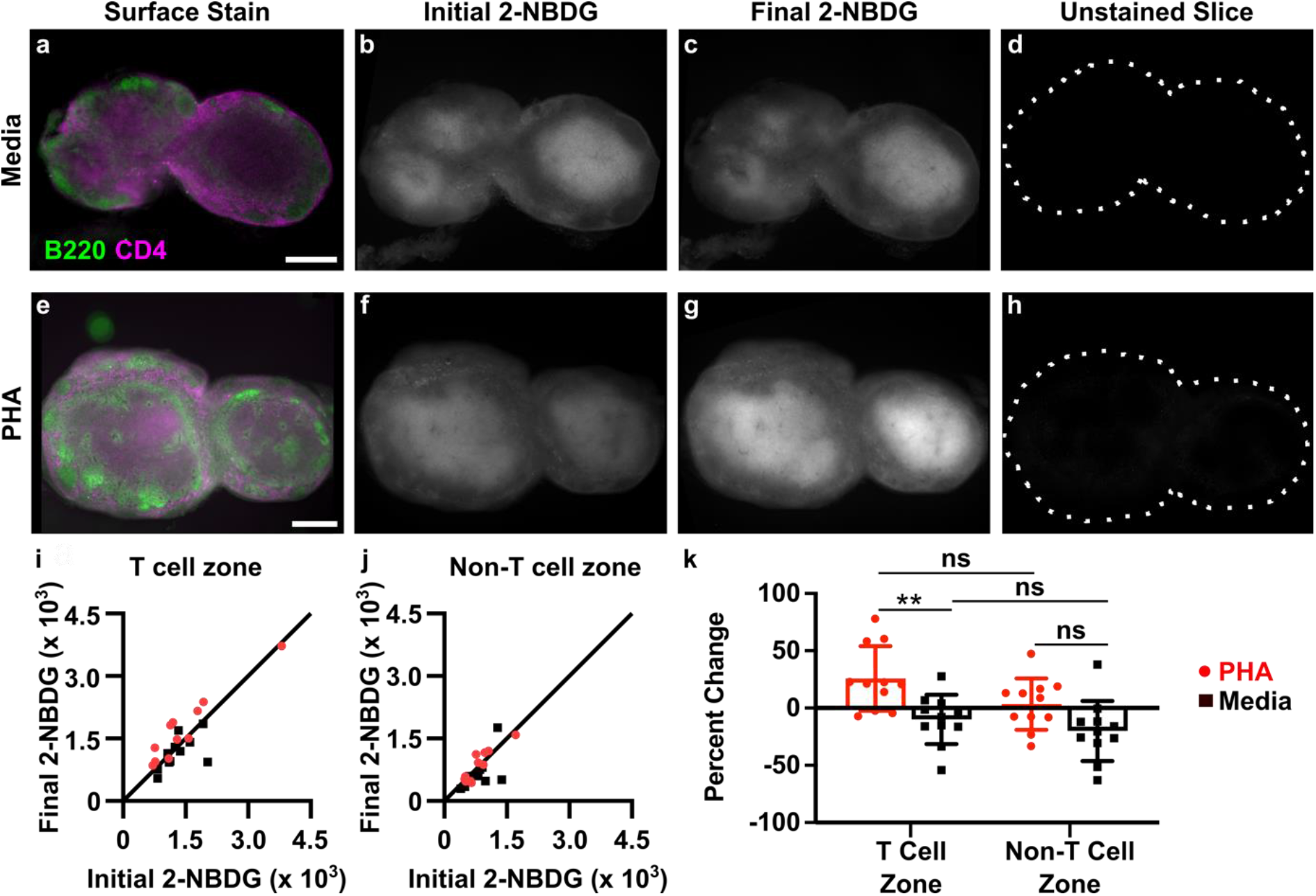
Spatially-resolved analysis of dynamic changes in glucose uptake in live tissues slices. (a-h) Representative images of lymph node slices cultured in media alone (a-d) or with 25 μg/ml PHA (e-h). (a,e) Immunofluorescence imaging to identify T cell zones, (b,f) initial 2-NBDG, (c,g) 2-NBDG following 3 hr culture, and (d,h) initial unstained image. All 2-NBDG images are scaled to the same brightness and contrast. Scale bars 500 μm. (i,j) Quantification of mean 2-NBDG intensity in the (i) T cell zones and (j) non-T cell zones, before and after 3-hr culture with PHA or media control. y=x line plotted for reference; dots on the line indicate no change in 2-NBDG uptake after culture. Each dot represents one slice. N=11 slices. (k) Data plotted as percent change from initial to final 2-NBDG intensity (0% = no change, 100% = doubling of intensity). Two-way ANOVA with Sidak’s multiple comparisons, ** represents p<0.01.

Ex vivo PHA stimulation produced a response that was detectable across the entire tissue on average (Fig. S3), and the response was localized specifically to the T cell zones (Fig. 5i-k). Quantitative image analysis showed a small increase in 2-NBDG uptake in the T cell zone after PHA stimulation (1.26-fold ± 0.27, mean ± stdev, N = 11 slices; Fig. 5i) that was significantly different than the response to media control (Fig. 5k). The magnitude of the response was consistent with cellular responses observed by flow cytometry in Figure 2, with variability characteristic of tissue slices. In contrast, locations outside of the T cell zone did not respond on average (1.03-fold ± 0.22, mean ± stdev, N = 11 slices; Fig. 5j) and were not significantly different than media controls (Fig. 5k). Notably, the media controls exhibited unchanged or even slightly diminished 2-NBDG uptake during the 3-hr culture (e.g., 0.90-fold ± 0.21 in T cell zones, mean ± stdev, N = 11 slices), indicating perhaps a small change in basal metabolic activity during culture of lymph node slices; this phenomenon remains to be further explored. In summary, these data demonstrate that the 2-NBDG assay can reveal changes in metabolic activity over time in live tissue with regional specificity, including the response to ex vivo stimulation.

## 4. CONCLUSIONS

We have developed a novel imaging-based assay that generates quantitative, spatially resolved maps of glucose uptake in live tissues at multiple time points during ex vivo culture. Compared to in vitro assays of glucose uptake, an ex vivo approach preserves the tissue microenvironment and provides insight into the organization of metabolically active cells. On the other hand, compared to in vivo glucose-sensing methods, ex vivo culture allows for more precise control over the timing and dose of stimulation in a particular organ. The assay described here is simple and robust, requiring only incubation and rinse steps with a commercially available fluorescent reagent, 2-NBDG. The 2-NBDG assay can be combined with live immunofluorescence labeling to determine which cell types internalize the most and least 2-NBDG. Furthermore, it can be used repeatedly to detect dynamics and stimulus-response behaviors in individual tissue slices. We anticipate that the simplicity of the assay will facilitate its adoption by other research laboratories for other types of soft tissue and models of disease.

When applied to live slices of lymph node tissue, this approach revealed varied glycolytic activity across the lymph node, which was clearly distinct from autofluorescence. The 2-NBDG signal was greatest in the paracortex (T cell zone), and ~70% of the signal could be attributed to CD4^+^ or CD8^+^ lymphocytes. The assay revealed that culture with a T cell mitogen triggered increased glucose uptake specifically in the paracortex, a pattern of glycolytic activity that could not have been detected with traditional analytical approaches. Future work may leverage this method to characterize region-specific responses in diseased tissues and evaluate the effects of therapeutic agents.

## ACKNOWLEDGEMENTS

We thank The Hartwell Foundation for generously funding this work and for helpful discussions on data analysis strategies. Research reported in this publication was supported by the University of Virginia 3Cavaliers Program through the Office of the Vice President for Research, and by the National Institute of Allergy and Infectious Diseases under Award Number R01AI131723 through the National Institutes of Health (NIH). The content is solely the responsibility of the authors and does not necessarily represent the official views of the National Institutes of Health. Some equipment for this project was funded by a Starter Grant Award from the Society of Analytical Chemists of Pittsburgh. DDD was supported by a Department of Chemistry Summer Scholarship and the College Science Scholars Program. The authors thank Ms. Alexis Chernish for experimental contributions at early stages of this project.

## Competing Interests Statement

The authors declare that they have no known competing financial interests or personal relationships that could have appeared to influence the work reported in this paper.

## Notes

### Competing Interest Statement

The authors have declared no competing interest.

### Summary of Updates

Updated manuscript.

