## Supplemental Information for "Spatially Resolved Measurement of Dynamic Glucose Uptake in Live Ex Vivo Tissues"

Electronic Supplementary Information for

Austin F. Dunn, Megan A. Catterton, Drake D. Dixon, Rebecca R. Pompano
University of Virginia, Department of Chemistry

### Supplemental Figures S1-S3

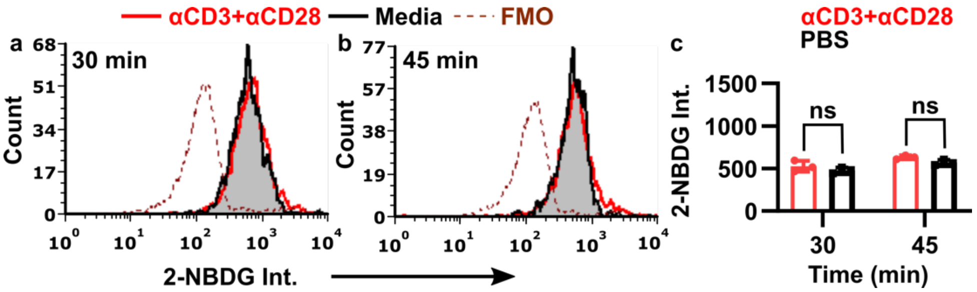


**Figure S1.** Used in a high-glucose diluent, 2-NBDG cannot differentiate stimulated from unstimulated lymphocytes. Here, cells were stimulated with anti-CD3+anti-CD28 (red) or left unstimulated (black), then cultured with 100 µM 2-NBDG in complete media for (a) 30 min and (b) 45 min. Samples treated and gated for T cells as in Fig. 2. (c) Median 2-NBDG intensity in T cells. Bars show mean ± std dev of n=3 replicates. Two-way ANOVA with Sidak's multiple comparisons, ns represents $p>0.05$.

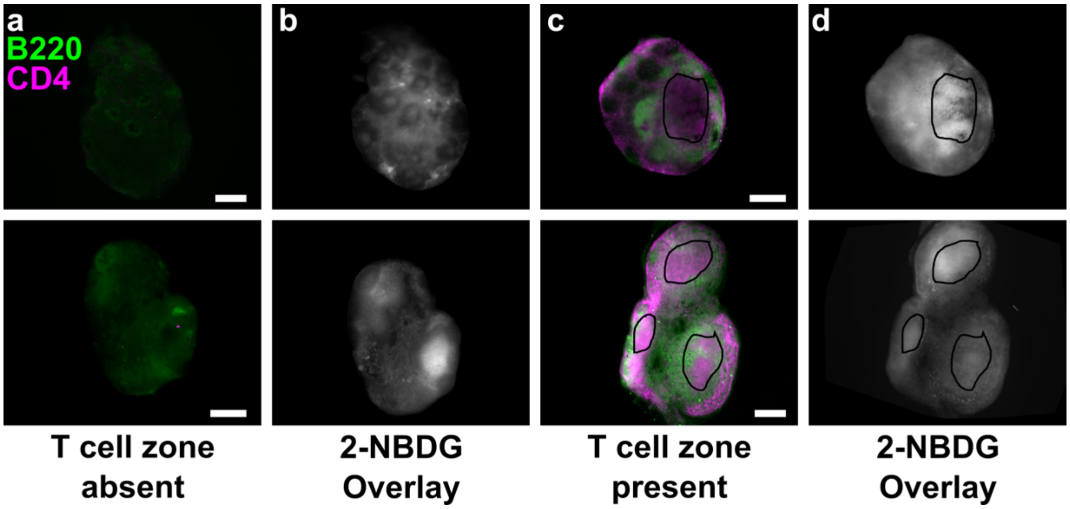


**Figure S2.** For the repeated-measures experiment (Figure 5), slices were excluded from analysis when T cell zones were absent or small. A blinded researcher outlined the nominal T cell zone(s) in each immunofluorescence image without having seen the 2-NBDG data. T cell zones were defined as $CD4^{high}$ $B220^{low}$ roughly circular or ovoid regions, usually towards the center of each lobe of the lymph node, though sometimes also near the edges. (a,b) Two representative slices, one per row, that were excluded from analysis due to the absense of any $CD4^{high}$ regions in the immunofluorescence images (a). 2-NBDG data from these tissues shown in (b). (c,d) Two representative slices that were included in analysis; designated "T cell zones" were outlined in the immunofluorescence images (c) and transposed onto 2-NBDG images in Fiji (d). Each fluorescent channel is leveled identically across all images. All scale bars 500 µm.

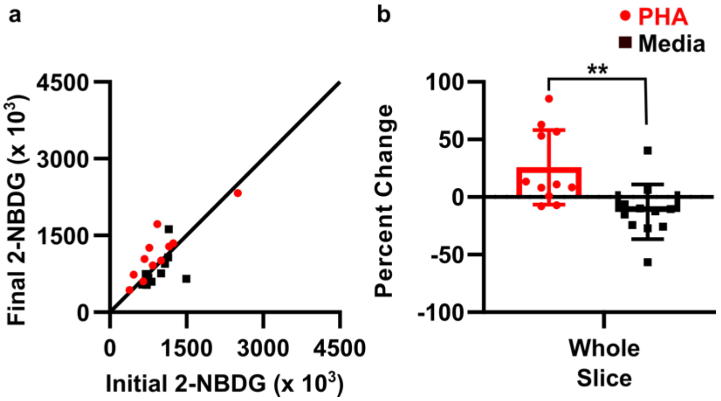


**Figure S3.** 2-NBDG intensity measured over the entire slice area defined by brightfield imaging. Same set of tissue slices as in Fig. 5. (a) Quantification of mean 2-NBDG intensity in the tissue slice, before and after 3-hr culture with PHA (red) or media control (black). y=x line plotted for reference; dots on the line show no change in 2-NBDG uptake after culture. Each dot represents one slice; N=11 slices. (b) Same data plotted as percent change from initial to final 2-NBDG intensity (0% = no change, 100% = doubling of intensity). Significance determined by Student's t-test. ** represents $p<0.01$.
